# Atom filtering algorithm and GPU-accelerated calculation of simulation atomic force microscopy images

**DOI:** 10.1101/2023.11.15.567294

**Authors:** Romain Amyot, Noriyuki Kodera, Holger Flechsig

## Abstract

Simulation atomic force microscopy computationally emulates experimental scanning of a biomolecular structure to produce topographic images that can be correlated with measured images. Its application to the enormous amount of available high-resolution structures, as well as to molecular dynamics modelling data, facilitates the quantitative interpretation of experimental observations by inferring atomistic information from resolution-limited measured topographies. The computation required to generate a simulated AFM image generally includes the calculation of contacts between the scanning tip and all atoms from the biomolecular structure. However, since only contacts with surface atoms are relevant, a filtering method shall highly improve the efficiency of simulation AFM computations. In this report we address this issue and present an elegant solution based on graphics processing unit (GPU) computations that significantly accelerates the computation of simulation AFM images. The method not only allows for visualization of biomolecular structures combined with ultra-fast synchronized calculation and graphical representation of corresponding simulated AFM images (live simulation AFM), but, as we demonstrate, can also reduce the computational effort during automatized fitting of atomistic structures into measured AFM topographies by orders of magnitude. Hence, the developed method will play an important role in post-experimental computational analysis involving simulation AFM. Implementation is realized in our BioAFMviewer software package for simulation AFM of biomolecular structures and dynamics.

## Introduction

Atomic force microscopy (AFM) can visualize the surface of biomolecular structures and high-speed AFM allows to follow conformational dynamics in real time under near physiological conditions [1-4]. A major limitation of AFM is that only morphological changes within the probed molecular surface can be detected, missing information required to fully explore functional mechanisms from imaging alone. Furthermore, since the scanning tip is too large to resolve structural detail, the interpretation of measured AFM topographic images including information below the sub nano-meter range is generally difficult.

Atomistic-resolution equilibrium molecular structures of most proteins are either known from a combination of experiments [5] or can be predicted by the artificial intelligence AlphaFold program [6]. On the other hand, functional conformational dynamics can be obtained from multi-scale molecular modelling [7-10]. The enormous amount of high-resolution protein data offers a great opportunity to better understand resolution-limited AFM scanning data. Simulation atomic force microscopy (S-AFM) is a computational method that mimics experimental tip-scanning of a biomolecule to transform its available atomistic structure into a simulated topographic image which can be compared with an experimental AFM image. A sufficiently high image similarity, quantified by correlation scores, indicates that the unknown atomistic arrangement behind a measured AFM topography may be best represented by the known template structure.

Various computational methods of systematically fitting atomistic-level or coarse-grained structural templates into AFM images have been developed [11-15] and evidenced to advance understanding of molecular processes beyond resolution-limited experimental imaging (see the recent review [16]). For example, the method of rigid-body fitting relies on exhaustive sampling molecular orientations of an atomistic template structure, each time applying S-AFM, until eventually the molecular arrangement whose S-AFM image has the highest image similarity with an experimental target is identified. The speed-limiting process during fitting is the computation of the S-AFM image of each sampled molecular orientation required for comparison to the target AFM image. This is because for a given molecular orientation, the computation generally involves evaluating contacts of the scanning tip with all atoms in order to generate a topographic image of a biomolecular structure. However, since the scanning tip can only contact the biomolecular surface without penetrating the structure, the computation can be significantly accelerated by filtering a set of relevant atoms. Here, we present a solution based on graphics processing unit (GPU) computations, combining the visualization of molecular structure with a filtering method conducted on-the-fly. The underlying idea is simple: For any arbitrary 3D molecular orientation of a biomolecular structure, the set of atoms which are directly exposed to the scanning tip and can potentially come into contact with it, contains exactly those which are displayed by rendering the corresponding graphical 2D view. We demonstrate the efficiency of this filtering method in applications of S-AFM computations for proteins of various shapes.

## Results

The S-AFM method typically rests on several approximations. The scanning tip is viewed as a rigid cone-shaped object with a probe sphere at its end, the molecular structure (sample) is represented by a static Van-der-Waals sphere atomic model placed on a solid surface (AFM substrate), and tip-sample interactions are considered as non-elastic collisions. Within such approximations S-AFM calculations are reduced to determine the single-point intersection of the 3D tip model with an atomic sphere, for which an explicit equation exists (from solving a second order polynomial equation, see [17] for details). However, computing the height topography of the entire sample for a given molecular orientation is still a time-consuming process. This is because for each cell along the scanning grid, generally contact points of the tip with all atoms have to be evaluated in order to determine the largest possible height value relative to the AFM substrate, roughly corresponding to heights measured in an AFM experiment. In fact, a ubiquitous situation is that the tip collides first with atoms that are beyond those belonging to the scanned grid cell. On the other side, it is obvious that the tip can only contact atoms which are exposed at the molecular surface. Therefore, the time required to compute a topographic image can in principle be reduced by introducing a method which, for a given molecular orientation, can filter surface atoms relevant for the S-AFM scanning process. The challenge is to construct a filtering algorithm that is fast enough to actually speed up the computation of the S-AFM image. For example, if filtering is more time consuming than the filter-free computation described above (for which just algebraic expressions must be evaluated), there is no efficiency gain.

The results are presented in the following way. We first explain our developed filtering method consisting of primitive filtering in the horizontal/vertical scanning direction and an elegant GPU-based solution to atom filtering in the lateral scanning direction. Then we explain the GPU-based computation of simulated AFM images considering the filtered subset of atoms. To evaluate the efficiency gain of the developed methods we apply simulation AFM computation to various proteins with different size and shape geometry and perform statistical analysis of the obtained data.

For the following explanation of the filtering method and simulation AFM computation, we consider the Van-der-Waals representation of a biomolecular structure in an arbitrary molecular orientation.

### Primitive X-Y filtering

We refer to primitive filtering as a selection method of relevant atoms in the horizontal and vertical (X, Y) scanning direction. For any arbitrary orientation of the biomolecular structure within the fixed screen canvas, first the tip scanning grid is obtained by determining the farthest possible tip contacts with the atomistic structure in the east and west directions (scanning range X), and those in the south and north directions (scanning range Y). We provide an illustration in Fig. 1. The size of grid cells corresponds to the given scan step. As a next step it would be reasonable to consider each individual grid cell with the tip placed at its center and calculate the tip contacts with all atoms to select the largest height value of the structure relative to the AFM substrate.

**Figure 1:**
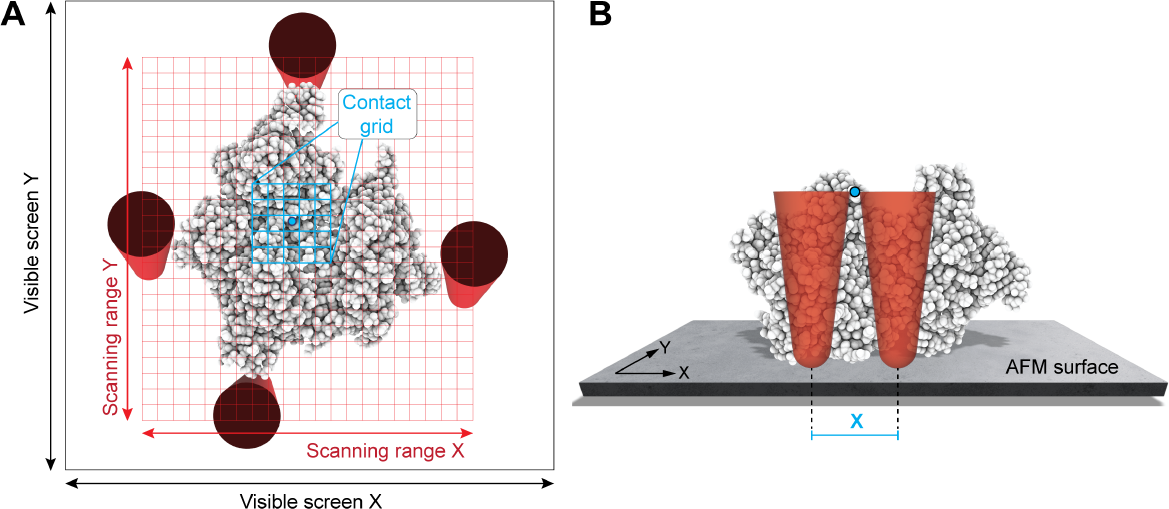
Primitive X-Y filtering. **(A)** The atomistic Van-der-Waals representation of a biomolecular structure is displayed in the scanning view perspective within the visible screen canvas (X, Y) outlined in black. The relevant scanning grid is shown in red, with the cell size corresponding to the given scan step. The four tip shapes illustrate the farthest possible tip contacts with the structure in the north, east, south, and west directions required to determine the scanning grid size. For a selected atom (blue circle), the computed tip contact grid is shown in blue. **(B)** The biomolecular structure placed on the AFM substrate in the front view perspective. For a selected atom (blue circle), the contact range of a cone-shaped scanning tip is illustrated for the X direction.

We replace this generally inefficient procedure by looping just once over all the atoms instead. For each atom a contact grid which covers the (X, Y) area from which the tip can touch this atom is determined as a subgrid of the scanning grid (Fig. 1). Its size apparently depends on the tip shape geometry and the distance of the atom to the AFM substrate. Then, for each cell of the contact grid the collision height of the tip with this atom is computed and compared with previous height values computed from a tip collision with other atoms, always storing the larger height value. Thus, after looping over all atoms each cell of the scanning grid is assigned a single value which represents the sample height relative to the AFM substrate as resulting from a convolution of the tip shape with the biomolecular surface. Simulation AFM using only primitive X-Y filtering (we refer to as the XY-F method) is obviously still highly inefficient. Nonetheless, we will later apply this method simply to demonstrate the supremacy of the more sophisticated filtering method and computation of simulation AFM described next.

### GPU-based lateral Z direction filtering of surface atoms

We developed an efficient method to filter the surface atoms in the lateral Z direction (i.e., in the vertical scanning direction of a biomolecular structure) based on graphics processing unit (GPU) computations. The underlying idea is that for any arbitrary 3D molecular orientation of a biomolecular structure, the set of atoms which are directly exposed to the scanning tip and can potentially come into contact with it, contains exactly those which are displayed by rendering the corresponding graphical 2D view.

To visualize objects on the computer screen graphic cards work with vertices and primitives. A primitive is the smallest unit used to build geometrical forms. Four main primitives are generally used: points, lines, triangles, and quads. To visualize atoms of a biomolecular structure as spheres (with corresponding Van-der-Waals radii) we use triangles as primitives. Spheres are therefore represented by a set of triangles, each of them having three vertices with attributes such as spatial position (x, y, z), orientation normal vector, color, etc. In a first step, the GPU processes all vertices with their attributes via the vertex shader (Fig. 2A). The vertex shader places the vertices relative to each other in the clip space, a GPU internal 3D coordinate system with coordinates ranging from −1 to 1 that can be viewed as a 3D version of what will become the visible screen. Vertices having clip coordinates beyond this range will be later clipped out of the scene. The vertex shader is programmable, and the programmer has full control of it. After processing all vertices, the GPU groups them to form the corresponding triangles in the clip space and then proceeds with rasterization.

**Figure 2:**
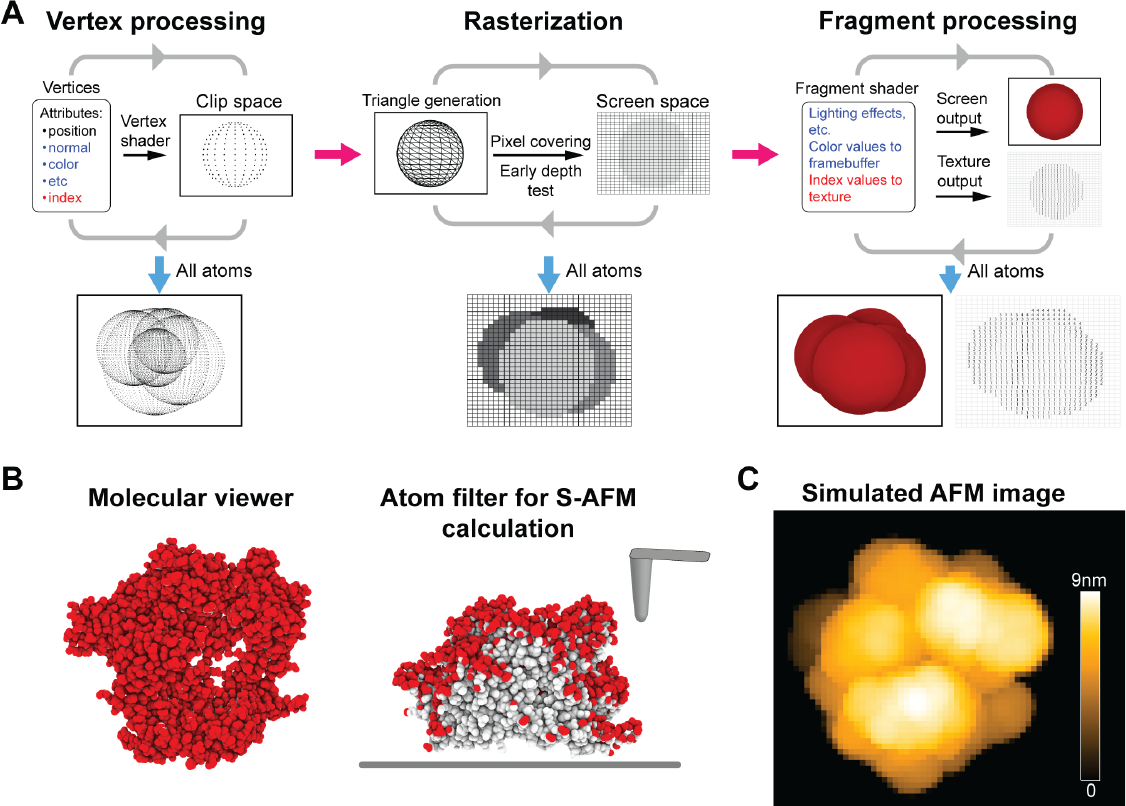
GPU workflow for lateral Z direction filtering of surface atoms. **(A)** Left, vertex processing: Every vertex is placed in the clip space coordinate system by the vertex shader. Its attributes are transferred to the next step. The attributes used in classical rendering are written in blue. The attribute used in our filtering method is written in red. Middle, rasterization: For each triplet of vertices, a triangle is formed, and fragments are generated based on the pixels covered by the triangle. The attributes of newly generated fragments result from the interpolation (or not) of the attributes of the three vertices. The depth test is applied to distinguish fragments in the front from others behind. Right, fragment processing: All fragments, which can now be viewed as pixels, enter the fragment shader. In classic rendering, the color is computed and rendered to the screen. In our filtering method, the index attribute of the pixel inherited from a vertex is written in a grid texture which can further be read on the CPU. **(B)** Left: Molecular structure of the Cas9 endonuclease protein (from PDB 4OO8) in the scanning view perspective showing the atoms that are probed by the AFM scanning tip. Right: Molecular structure in the front view perspective. The AFM substrate surface is indicated by the thick gray line. The surface atoms remaining after filtering (∼24% of total atoms) are shown in red color. **(C)** Simulated AFM image for the scanning view in (B) computed from the set of filtered atoms.

Rasterization is the process by which the GPU transforms the 3D clip space into what will become the visible screen space, i.e., a 2D canvas of pixels (Fig. 2A). Rasterization consists of determining which pixels are covered by a particular triangle and generates so called fragments accordingly. A fragment is characterized by the 2D position of the pixel it belongs to plus the attributes inherited from the vertices of the triangle including the interpolated clip space z-coordinate interpreted as the depth value of the fragment by the GPU. During processing all triangles, many fragments will obviously share the same pixel position. However, an early depth test allows to compare the depth of a newly generated fragment with that of the current fragment for a given pixel position keeping only the least deep fragment. In such a situation the result of the rasterization process is a pixel grid which consists of filtered fragments, each fragment representing the least deep position for a given pixel.

Finally, all remaining fragments from the previous rasterization process undergo fragment shading to determine a color for the processed fragment (Fig. 2A). The normal and color attributes are usually used to compute light effects and the 4D output vector from the fragment shader (RGB values plus opacity) will be interpreted by the GPU as a color for a particular pixel. In the context of visualizing a biomolecular structure, completed fragment shading would result in rendering the Van-der-Waals representation on the computer screen (*render to screen* pathway). I.e., for a given molecular orientation only the visible atoms are displayed and separated from the hidden non-visible atoms. Obviously, this separation would exactly correspond to filtering the subset of atoms that would be accessible to the scanning tip in that particular 3D orientation. Since similar to the vertex shader the fragment shader is programmable, we are able to implement the desired atom filter algorithm for S-AFM. This is possible via by-passing the rendering process and directing the fragment shader output to a texture (*render to texture* pathway), which can be viewed as an off-screen grid of pixels that can be read on the central processing unit (CPU). Therefore, the atom filtering procedure for an arbitrary orientation of the biomolecular structure to facilitate computation of the corresponding S-AFM image can be performed independent from visualizing the 3D molecular structure (Fig. 2B).

Like in the XY-F method, simulation AFM calculations can now be performed using the largely reduced set of filtered atoms to obtain the simulation AFM image (Fig. 2C). We refer to this method as the XYZ-F method and evaluate its performance against the XY-F method and the ultimate method described next.

### GPU-based computation of simulation AFM images

Finally, we developed a method which takes into account the filtered set of atoms to efficiently compute the simulation AFM image employing an adaptation of the GPU workflow described above.

The lateral Z direction filtering uses the normal rendering process of the graphic card to output the indices of visible atoms. The simulation AFM calculation method we developed manipulates the workflow of the GPU from its original purpose to let the GPU parallelize the calculations with a minimal coding effort and without having to handle problems related to parallel computations. I.e., there is no need to code a complicated program for parallelization and to take care of all related problems (such as concurrency access).

For this method, each atom is represented by a point primitive with the 3D position and corresponding Van-der-Waals radius as attributes. The rendering texture corresponds to the scanning grid (Fig. 1A, red grid) in which each cell represents one pixel. The vertex shader operates in the same way as during filtering, i.e., vertices (in this case points) are mapped onto the clip space. Before the rasterization, vertices are intercepted by the geometry shader. The geometry shader is an optional programmable shader which takes a primitive as input and outputs one or more primitives (which can be of different type from the input primitive). For our purpose, we employ the geometry shader to generate for each filtered atom the corresponding tip contact sub-grid (Fig. 1A, blue grid). I.e., for each point primitive as the input the shader generates as an output a quad primitive with the size and position of the tip contact sub-grid for the corresponding atom. During the rasterization process the obtained quads are mapped onto the pixel screen space and fragments are generated. Each fragment encodes the (X, Y) pixel position and the attributes of the point primitive it originated from, i.e., the corresponding atom position (x, y, z) and Van-der-Waals radius. The early depth test is inactivated such as all generated fragments resulting from all previously filtered atoms will undergo the fragment shader. Generally, many fragments with different attributes share the same pixel position. The fragment shader is used to compute for each fragment the contact height of the tip (located over the corresponding pixel position) and the corresponding atom with respect to the AFM substrate surface, and the obtained height values are stored in the texture. The depth test takes place at this stage keeping for each pixel of the texture only the fragment with the largest height value. Hence, the resulting texture contains for each pixel the largest height value obtained from simulating scanning of the atomistic biomolecular structure in an arbitrary 3D orientation relative to the fixed AFM surface with a cone-shaped tip over the scanning grid with a prescribed spacing according to the experimental scan step. The corresponding simulation AFM image is visualized by mapping height values to a given color scale and coloring all pixels accordingly (Fig. 2C).

### Quantification of computational efficiency gain

To quantify the efficiency gain of the developed atom filtering methods and GPU calculations we considered exhaustive sampling, which is a procedure employed during fitting of structural data into AFM topographic images. This method samples rigid-body orientations of a given structural template in 3D space, each time computing the corresponding simulation AFM image which is compared with an experimental target image. As structural templates we used proteins which differ in size and shape:

- the hepatitis B virus capsid cryo-EM ball-shaped structure with a diameter of ∼36 nm and a total of 270,960 atoms (constructed from PDB 6BVF)
- a model of the actin filament structure containing 24 subunits and having a rod-shape with ∼10 nm diameter, ∼70 nm length, and a total of 70,464 atoms (from Ref. [18])
- the rotor-less F_1_-ATPase motor cube-like structure with lengths of ∼10 and ∼12.5 nm and a total of 21,867 atoms (PDB 1SKY)
- the small global-shaped relaxin protein with lengths of ∼3 nm and ∼5 nm having just 755 atoms (PDB 6RLX)

The molecular structures are shown in Fig. 3A. For the four different protein structures rigid-body orientations were sampled uniformly in 3D space along a grid with a 5 degree spacing, resulting in (360/5)^3^=373,248 conformations that were placed on a fixed plate and scanned in the lateral direction by the scanning tip across the XY scanning grid. For each orientation we then recorded the computation time required to execute the filtering method and to calculate the corresponding simulated image (without visualizing it). We considered the three filtering methods described before, i.e., primitive XY filtering (XY-F), filtering including the lateral scanning direction Z (XYZ-F), and the latter with GPU calculations (XYZ-F2). The corresponding distributions of computation times for the four proteins are shown in Fig. 3A and discussed below.

**Figure 3:**
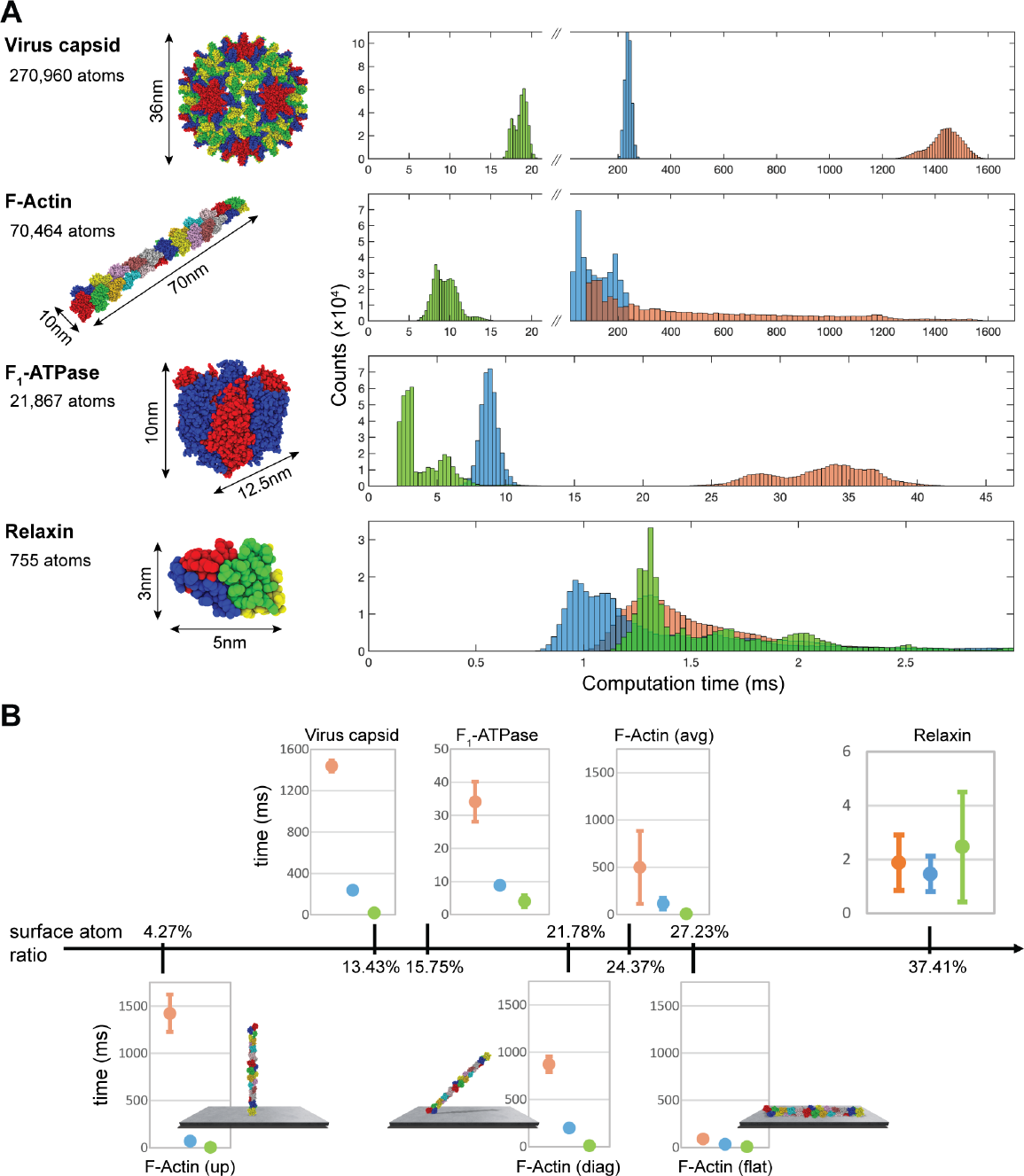
Quantification of computational efficiency gain by atom filtering methods. **(A)** Molecular structures of the investigated proteins are shown in the Van-der-Waals representation on the left side. The number of atoms is given, and their sizes are indicated. For each protein distributions of the computation time required to execute the filtering method and to calculate the simulation AFM image for molecular orientations obtained from exhaustive sampling are shown as histograms for the XY-F (orange color), XYZ-F (blue color), and XYZ-F2 (green color) atom filtering methods on the right side. **(B)** The same results are represented employing the surface atom ratio as a coordinate. For each filtering method the data is shown as mean value and standard deviation of its computation time distribution (solid circle and error bar). For the upper row data (virus capsid, F_1_-ATPase, F-Actin (avg), relaxin) the surface atom radio was obtained as an average over all considered molecular orientations. The bottom row data is for the actin filament considered in the upright, diagonal, and flat orientations with respect to the AFM substrate surface. For all plots of F-actin data the same scale was used for better comparison.

For the virus capsid structure, the calculation of an S-AFM image by the XYZ-F method (238 ±12 ms) is faster than the XY-F method (1438 ± 61 ms) by a factor larger than 5 and is further enhanced by the XYZ-F2 method (19 ± 0.8 ms) by a factor larger than 10. To put those numbers into perspective, this means that the computation of S-AFM images during rigid-body fitting of the capsid structure into an actual AFM image (with our chosen parameters and hardware setup) would on the average take ∼149 hours (i.e., 6.2 days) using the inefficient XY-F methods, versus only ∼2 hours required with the XYZ-F2 implementation.

For the F-actin structure the computation time of an S-AFM image by the XY-F method shows a very wide distribution (499 ± 386 ms) which is due to the special rod-shape of the molecule. For upright orientations with respect to the AFM substrate the XY contact grid of the tip for each atom is generally large because most atoms are located far from the substrate, hence resulting in a large number of tip-atom collisions to be calculated. In contrast, for orientations placed flat on the substrate the XY contact grids of atoms have much smaller size and computation is therefore faster. In such cases speed improvements by additional Z-filtering are less relevant - distributions of the XY-F and XYZ-F methods even overlap - whereas for filament positions oriented towards the upright shape Z-filtering becomes increasingly important and the XYZ-F method significantly enhances the computation speed as shown by the time distribution (117 ± 63 ms). The XYZ-F2 method supersedes others by improving the computation speed further by one order of magnitude (9.5 ± 1.5 ms).

For the relatively small F_1_-ATPase protein the calculation of a simulated AFM image by the XYZ-F method (8.8 ± 0.7 ms) is faster than the XY-F method (34 ± 6 ms) by a factor of ∼4 and is further enhanced by the XYZ-F2 method (4 ± 2 ms) by a factor of ∼2. This means that the computation of S-AFM images during rigid-body fitting of the F_1_ structure into an actual AFM image (with our chosen parameters and hardware setup) would on the average take 212 minutes, versus only 25 minutes required with the XYZ-F2 implementation.

Results of computation times obtained for the relaxin protein, i.e., (1.88 ± 1.03 ms) for the XY-F method, (1.46 ± 0.66 ms) for the XYZ-F method, and (2.47 ± 2.04 ms) for the XYZ-F2 method, clearly demonstrate the irrelevance of atom filtering for simulation AFM of very small molecular structures.

The numerical investigations lead us to the following conclusions. Generally, our developed atom filtering methods significantly speed up the calculation of simulation AFM images. The efficiency gain depends on the size and shape of proteins and is apparently larger for cases in which the number of surface atoms is small as compared to the total number of atoms. To clearly illustrate this aspect, we represent in Fig. 3B the results discussed above employing the ratio of surface atoms and all protein atoms as a coordinate. We emphasize again that for very large protein structures the calculation time of a simulated AFM image can be reduced by ∼2 orders of magnitude by the application of the developed filtering method. While for proteins with typical sizes of ∼10 nm a significant speed up in calculation can be realized, the filtering methods become irrelevant for very small proteins with a generally larger surface atom ratio.

## Summary

Post-experimental analysis of biomolecular dynamics visualized by AFM experiments employing computational methods plays an increasingly important role. In this situation, simulation atomic force microscopy is the cornerstone method allowing to correlate atomistic resolution biomolecular data with resolution-limited measured topographies.

Application of simulation AFM allows to employ the enormous amount of available structural data, as well as data obtained from molecular dynamics simulations, to facilitate the interpretation of resolution-limited imaging towards an atomistic level understanding of measured nanoscale processes. E.g., rigid-body fitting by exhaustive sampling molecular orientations of a structural template [13,15], or flexible fitting methods resolving conformational changes [11,12,14], are based on the execution of simulation AFM in each iteration step towards finding the atomistic conformation that best matches with a target experimental AFM image. As we demonstrate, our developed method based on the GPU workflow can accelerate the computation of simulation AFM images by orders of magnitude, which can make a difference between several days versus a couple of hours required for fitting. Hence, this method will play an important role in computational analysis involving calculations of simulation AFM.

We postulate that our developed method of atom filtering and GPU-acceleration of simulation AFM images presents the algorithm of ultimate efficiency and further efficiency gain can only be achieved by exploiting faster hardware.

The developed method is implemented in our BioAFMviewer software package for simulation AFM of biomolecular structures and dynamics [17,19], where is allows for visualization of biomolecular structures (of potentially massive size) combined with ultra-fast synchronized calculation and graphical representation of corresponding simulated AFM images (live simulation AFM).

## Author contributions

R.A.: developed methods, developed software, analyzed data, discussed results, produced figures, wrote paper (revision); N.K.: supervised project, wrote paper (revision); H.F.: supervised project, analyzed data, discussed results, produced figures, wrote paper (initial and revision);

## Data availability

For the implementation of our method we used the OpenGL library [20,21]. Programming code to implement the developed methods is available upon request (to R.A. or H.F.). After accepted publication data will be made available in an open access repository.

